# Physiological changes throughout the ear due to age and noise - a longitudinal study

**DOI:** 10.1101/2021.11.25.470017

**Authors:** Alix Blockley, Daisy Ogle, Charlie Woodrow, Fernando Montealegre-Z, Ben Warren

**Author notes:** **Corresponding author: Ben Warren,**. these authors contributed equally (Alix Blockley, Daisy Ogle).

## Abstract

Biological and mechanical systems, whether by their overuse or their aging, will inevitably fail. Hearing provides a poignant example of this with noise-induced and age-related hearing loss. Hearing loss is not unique to humans, however, and is experienced by all animals in the face of wild and eclectic differences in ear morphology and operation. Here we exploited the high throughput and accessible tympanal ear of the desert locust, *Schistocerca gregaria* (mixed sex) to rigorously quantify changes in the auditory system due to noise exposure (3 kHz pure tone at 126 dB SPL) and age. We analysed tympanal dispalcements, morphology of the auditory Müller’s organ and measured activity of the auditory nerve, the transduction current and electrophysiological properties of individual auditory receptors. We found that noise mildly and transiently changes tympanal displacements, decreases both the width of the auditory nerve and the transduction current recorded from individual auditory neurons. Whereas age – but not noise - decreases the number of auditory neurons and increases their resting potential. Multiple other properties of Müller’s organ were unaffected by either age or noise including: the number of supporting cells in Müller’s organ or the nerve, membrane resistance and capacitance of the auditory neurons. The sound-evoked activity of the auditory nerve decreased as a function of age and this decrease was exacerbated by noise, with the largest difference during the middle of their life span. This ‘middle-aged deafness’ pattern of hearing loss mirrors that found for humans exposed to noise early in their life.

**Key point summary:**

- Age and noise lead to hearing loss.
- Tympanal displacements have a transient and delayed reduction after noise.
- The number of auditory neurons decreases with age. The width of the auditory nerve is reduced with noise exposure.
- Locusts repeatedly exposed to noise lost their hearing compared to silent controls, but their hearing became similar to old silent control locusts due to age-related hearing loss dominating.
- The electrophysiological properties of the auditory neurons and the transduction current remained unchanged for aged locusts but repeated noise exposure led to a cumulative decrease in the transduction current.

## Introduction: The multifaceted causes and consequences that lead to hearing loss

Nothing lasts forever. Left unchecked, systems inevitably fail. Two of the biggest causes are age and their use. Ears are a fascinating and relevant biological example of this principle. Age-related hearing loss is something that we will inevitably experience and noise-induced hearing loss is responsible for an estimated third of all worldwide hearing loss (Bethesda, 1990). Audiometric measurements from workers in consistently noisy environments – before the enforcement of ear protection - has gifted us a crucial quantitative and longitudinal dataset to understand the interaction of noise and age on hearing loss (Burns and Robinson, 1970; Passchier-Vermeer, 1968, 1977). We know that threshold shifts due to noise can, with a small corrective factor (Corso, 1980), be simply added to threshold shifts due to noise (Macrae, 1971; Miils, 1998ILR; Corso et al., 1976; Mollica, 1969). From this dataset Corso (1980) found that noise-exposed individuals differed most from their silent counterparts during the middle of their life but are very similar in older life. This is because age-related hearing loss dominates in later life. This simplistic mathematical interaction of age and noise on hearing thresholds is in stark contrast to the complex and insidious physiological deficits found in ears across animal phyla responsible for hearing loss.

The physiological and anatomical changes that co-occur with hearing loss present at multiple levels of the auditory system. This stretches from the middle ear and tympanum (Etholm and Belal, 1974; Ruah et al., 1991; Rolvien et al., 2018), to the inner ear supporting cells (Thorne and Gavin 1984; Shi & Nuttall, 2003) and auditory receptors (Keithley and Feldman, 1982; Liberman and Beil, 1978; Bohne and Harding, 2000; Wu et al., 2019; Jeng et al., 2020), including their synapses to the auditory nerve (Kujawa and Liberman, 2009, 2015; Wu et al., 2019) and the central nervous system where sound is processed (Fetoni et al., 2013). The most profound and best quantified change in the auditory system is loss of hair cells (Coleman, 1976; Keithley and Feldman, 1982; Bhattacharyya and Dayal, 1985; Dayal and Bhattacharyya, 1986; Li and Hultcrantz, 1994; Dixon and Ward, 1971; Pinheiro et al., 1973; Clark et al., 1974; Moody et al., 1978). Loss of hair cells can lead to loss of the spiral ganglion neurons onto which they synapse (Dupont et al., 1993; McFadden et al., 2004; Bohne and Harding, 2000; Johnsson and Hawkins, 1972). The auditory nerve formed by the axons of the spiral ganglion neurons and Schwann cell folds decreases in thickness after noise exposure (Pilati et al., 2012; Tagoe et al., 2014; Wan and Corfas, 2017). The complex correlation of multiple physiological deficits with hearing loss appears to have mistakenly led to the cause being assumed to the most consistent deficit – for instance with age-related hearing loss being due to the most common stria vascularis pathology (Ramadan and Schuknecht, 1989; Wu et al., 2020). In short, we lack the experiments and quantitative data to track the progression of known changes in the auditory system and weigh up their contribution to hearing loss

Adding to this complexity, it is not known which physiological deficits are caused by noise, age or a combination of both. There are some breakthrough studies that have proportioned specific deficits into noise or age (Kujawa and Liberman, 2006, 2009; Fernandez et al., 2015). Kujawa and Liberman (2009) found that ribbon synapses – those between hair cells and spiral ganglion neurons – sharply decrease after noise but appear to be only mildly decreased with age. And Wu et al. (2020) found that most hearing loss can be explained by hair cell loss not stria vascularis degeneration.

At present high-powered quantification of multiple physiological deficits in one model organism in response to noise or age over the course of its lifespan are lacking. Further to this, age-related hearing loss and noise-induced hearing loss are thought to be very distinct processes (Yang et al., 2015), which could be independent of each other (Corso et al., 1976). In this work we rigorously quantify the physiological changes responsible for hearing loss – found across animal phyla - over the lifespan of the desert locust, *Schistocerca gregaria*. Using a high-powered approach we measure: displacements *in vivo* of the tympanum, the morphology and anatomy of the auditory Müller’s organ, electrophysiological properties of the auditory receptors and their transduction currents *ex vivo* and sound-induced auditory nerve responses *in vivo*.

## Materials and methods

### Locust husbandry

Desert locusts (*Schistocerca gregaria*) were reared in crowded conditions (phase gregaria) on a 12:12 h light:dark cycle at 36:25°C. Locusts were fed on a combination of fresh wheat and bran *ab libitum*. We used nearly equal numbers of female and male *Schistocerca gregaria* from the Leicester ‘Locust Labs’ laboratory stock. The cuticle of locusts continues to sclerotize up until they change (yellow for males, grey for females) colour at ∼ 10 days post their last moult. And recordings from single auditory neurons are not possible until this time. Therefore we used locusts between 10 and 34 days post imaginal moult. In all experiments day 1 corresponds to locusts 10 days post their last moult. We used new locusts for each set of experiments. We are willing and open to share our locust strains with other research groups.

### Survival assay and life span in locusts

We counted the deaths of locusts over the 24 days (Fig. 1) whilst subjecting the locusts to identical conditions as the experimental locusts. Approximately 50% of locusts were dead at 12 days post first noise exposure or mock noise exposure (for control group) and there was no stark difference between noise-exposed and control locusts. After this time the rate of locust mortality decreased drastically so that very few locusts died over the next 12 days until we stopped measuring.

**Fig. 1.**
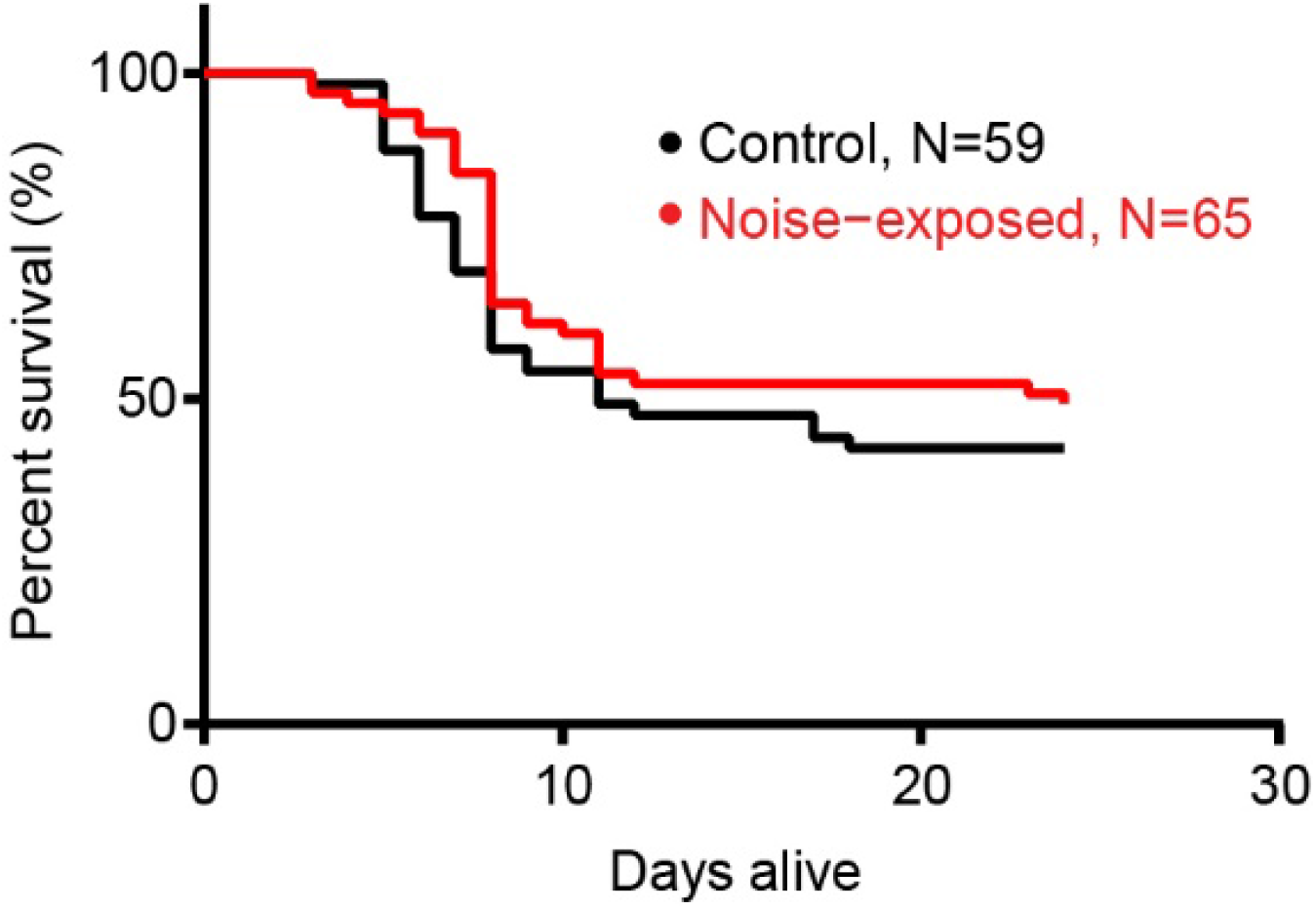
Survival of control and noise-exposed locusts. Day 1 is approximately 10 days post their last moult into adults.

### Noise exposure and acoustic stimulation

The wings of all locusts (control and noise-exposed) were cut off at their base to increase noise exposure of the conditioning tone to their tympanal ears, which are otherwise covered by their wings. Between ten and twenty locusts, for both the noise-exposed group and the control group, were placed in a cylindrical wire mesh cage (8 cm diameter, 11 cm height). Both cages were placed directly under a speaker (Visaton FR 10 HM 4 OHM, RS Components Ltd). For the noise-exposed group only, the speaker was driven by a function generator (Thurlby Thandar Instruments TG550, RS Components Ltd) and a sound amplifier (Monacor PA-702, Insight Direct Ltd) to produce a 3 kHz tone at 126 dB SPL, measured at the top of the cage where locusts tended to accumulate. Throughout the paper we refer to noise that the locusts are exposed to as this 3 kHz 126 dB SPL pure tone. This tone was played continuously for 12 hours overnight (21:00-09:00) for the noise-exposed group during their natural darkness period. The control group was housed in an identical cage with a silent speaker for 12 hours. All recordings were performed within a twelve hour window during the day. Sound Pressure Levels (SPLs) were measured with a microphone (Pre-03 Audiomatica, DBS Audio) and amplifier (Clio Pre 01 preamp, DBS Audio). The microphone was calibrated with a B&K Sound Level Calibrator (CAL73, Mouser Electronics). For laser measurements, hook electrode and patch-clamp recordings, the locust ear was stimulated with the same speaker and amplifier as above with a 3 kHz pure tone duration of 0.5s. For hook electrode recordings the 3 kHz tone had a rise and fall time of 2 ms. Tones were played three times for each locust at each SPL and the average response taken for each SPL. For intracellular recordings from individual auditory neurons the speaker was driven by a custom made amplifier controlled by an EPC10-USB patch-clamp amplifier (HEKA-Elektronik) controlled by the program Patchmaster (version 2×90.2, HEKA-Elektronik) running under Microsoft Windows (version 10).

### Biomechanical measurements of the tympanum with laser Doppler vibrometry

For *in vivo* measurements of the tympanum, locusts were mounted in natural dorso-ventral orientation following removal of wings and hind legs, and fixed to a copper platform using blue tac (Bostik, Colombes, France). They were positioned so that their tympanum was perpendicular to the micro-scanning Laser Doppler Vibrometer (PSV 500 with close up unit and 150 mm lens, Polytec, Waldbronn, Germany). The stimulus was produced with a waveform generator (SDG 1020, Siglent, China), and delivered via a stereo amplifier (SA1 power amplifier, Tucker-Davis Technologies, Alacchua, Florida) to a loudspeaker (MF1, Tucker-Davis Technologies, Alacchua, Florida) positioned 15 cm away from the animal to avoid nearfield measurement. The stimulus amplitude was obtained by manually changing the voltage within the waveform generator, and recording the SPL using a 1/8” microphone (Type 4138, Brüel & Kjaer, Germany) with built in preampflifer (B&K 2670, Brüel & Kjær, Denmark), calibrated using a sound-level calibrator (Type 4237, Brüel & Kjaer, Denmark), via a conditioning amplifier (Nexus 2690-OS1, Brüel & Kjær, Denmark). Displacement data of tympanum vibration was collected using the PSV internal data acquisition board with a 128 ms sample length at a sampling rate of 512,000 Hz, averaged 10 times per sample. Experiments were carried out in an acoustic booth (IAC Acoustics, Series 120a, internal dimensions of 2.8 m x 2.7 m x 2 m) on a pneumatic vibration isolation table (Nexus Breadboard (B120150B), 1. 2 m x 1.5 m x 0.11 m,, Thor Labs, USA)). Displacements were measured as the average peak-to-peak displacement.

### In vivo hook electrode recordings from auditory nerve six

Locusts were secured ventral side up in plasticine. A section of the second and third ventral thoracic segment was cut with a fine razor blade and removed with fine forceps. Tracheal air sacks were removed to expose nerve six and the metathoracic ganglia. Hook electrodes constructed from silver wire 18 μm diameter (AG549311, Advent Research Materials Ltd) were hooked under the nerve and the nerve was lifted out of the haemolymph. A mixture of 70% Vaseline and 30% paraffin oil was applied through a syringe to coat the auditory nerve to stop it drying out. Signals were amplified 1,000 times by a differential amplifier (Neurolog System) then filtered with a 500 Hz high pass filter and a 50 kHz low pass filter. This amplified and filtered data was sampled at 25 kHz by Spike2 (version 8) software running on Windows (version 10). To quantify the compound spiking activity of the auditory nerve we used Matlab (Version R2020a, Mathworks Inc.) and rectified the nerve signal and integrated the area underneath. We computed this for the 0.5 s of sound-evoked neural activity and for 60 s background nerve activity before the tones and the background activity between the tones. To compute the σ ratio we divided the sound-evoked response by the background neural activity. N.B. the locust treatment was blinded to the experimenter until all data was collected and analysed.

### Dissection of Müller’s Organ and isolation of Group-III auditory neurons

Whole cell patch clamp recordings per performed on group-III auditory neurons because they form the majority of auditory neurons of Müller’s organ (∼46 out of ∼80) (Jacobs et al., 1999), they are the most sensitive auditory neurons of Müller’s organ (Römer, 1976) and are broadly tuned to the 3 kHz we used for noise-exposure (Warren and Matheson, 2018). For intracellular patch-clamp recordings from individual auditory neurons the abdominal ear, including Müller’s Organ attached to the internal side of the tympanum, was excised from the first abdominal segment, by cutting around the small rim of cuticle surrounding the tympanum with a fine razor blade. Trachea and the auditory nerve (Nerve 6) were cut with fine scissors (5200-00, Fine Science Tools), and the trachea and connective tissue removed with fine forceps. This preparation allowed perfusion of saline to the internal side of the tympanum, necessary for water-immersion optics for visualizing Müller’s Organ and the auditory neurons to be patch-clamped, and concurrent acoustic stimulation to the dry external side of the tympanum. The inside of the tympanum including Müller’s Organ was constantly perfused in extracellular saline.

To expose Group-III auditory neurons for patch-clamp recordings, a solution of collagenase (0.5 mg/ml) and hyaluronidase (0.5 mg/ml) (C5138, H2126, Sigma Aldrich) in extracellular saline was applied onto the medial-dorsal border of Müller’s Organ through a wide (12 μm) patch pipette to digest the capsule enclosing Müller’s Organ and the Schwann cells surrounding the auditory neurons. Gentle suction was used through the same pipette to remove the softened material and expose the membrane of Group-III auditory neurons. The somata were visualized with a Cerna mini microscope (SFM2, Thor Labs), equipped with infrared LED light source and a water immersion objective (NIR Apo, 40x, 0.8 numerical aperture, 3.5 mm working distance, Nikon) and multiple other custom modifications. For a full breakdown of the microscope components and how to construct a custom patch-clamp microscope for ∼£12k see: https://www2.le.ac.uk/departments/npb/people/bw120.

### Electrophysiological recordings and isolation of the transduction current

Electrodes with tip resistances between 3 and 4 MΩ were fashioned from borosilicate class (0.86 mm inner diameter, 1.5 mm outer diameter; GB150-8P, Science Products GmbH) with a vertical pipette puller (PC-100, Narishige). Recording pipettes were filled with intracellular saline containing the following (in mM): 170 K-aspartate, 4 NaCl, 2 MgCl2, 1 CaCl2, 10 HEPES, 10 EGTA. 20 TEACl. Intracellular tetraethylammonium chloride (TEA) was used to block K^+^ channels necessary for isolation the transduction. To further isolate nad increase the transduction current we also blocked spikes with 90 nM Tetrodotoxin (TTX) in the extracellular saline. During experiments, Müller’s Organs were perfused constantly with extracellular saline containing the following in mM: 185 NaCl, 10 KCl, 2 MgCl2, 2 CaCl2, 10 HEPES, 10 Trehalose, 10 Glucose. The saline was adjusted to pH 7.2 using NaOH. The osmolality of the intracellular and extracellular salines’ were 417 and 432 mOsm, respectively.

Whole-cell voltage-clamp recordings were performed with an EPC10-USB patch-clamp amplifier (HEKA-Elektronik) controlled by the program Patchmaster (version 2×90.2, HEKA-Elektronik) running under Microsoft Windows (version 7). Electrophysiological data were sampled at 50 kHz. Voltage-clamp recordings were low-pass filtered at 2.9 kHz with a four-pole Bessel filter. Compensation of the offset potential were performed using the “automatic mode” of the EPC10 amplifier and the capacitive current was compensated manually. The calculated liquid junction potential between the intracellular and extracellular solutions was also compensated (15.6 mV; calculated with Patcher’s-PowerTools plug-in from www3.mpibpc.mpg.de/groups/neher/index.php?page=software). Series resistance was compensated at 77% with a time constant of 100 μs.

### Staining and confocal microscopy

Locusts were secured ventral side up in plasticine. A square section of the second and third ventral thoracic segment was cut with a fine razor blade and removed with fine forceps and set aside. Tracheal air sacks were removed to expose nerve six and the metathoracic ganglia. The thoracic cavity was filled with locust extracellular saline and auditory nerve six was pinched and broken at the top of the nerve, close to the metathoracic ganglion with fine forceps and the cut end of nerve six placed on the thoracic cuticle outside the thoracic cavity at the posterior end. A well was formed around the nerve end with petroleum jelly (Vaseline, Boots) using a syringe (1.1mm diameter). The well was filled with Neurobiotin (5% m/v, SP-1120, Vector Laboratories) dissolved in distilled water before a lid was fashioned with more petroleum to seal the well. The square of thoracic cuticle was replaced onto the locust thoracic cavity and sealed back in place with petroleum jelly, to prevent desiccation. Locusts were incubated overnight at 4°C to allow Neurobiotin to diffuse along the nerve and fill the auditory neurons of Müller’s organ. Following overnight incubation, whole ears were excised from the first abdominal segment, as described above. Whole locust ears were fixed in 4% paraformaldehyde (P6148, Sigma Aldrich) dissolved in Phosphate Buffer Saline (PBS) for 24 hours at 4°C, in a single well of a 24 plastic-well plate. Following fixation, ears were washed in PBS (3 × 10 minutes) at room temperature, before being stored in PBS at 4°C. Locust ears were washed in PBS, with 0.2 % m/v Triton-X100 and 5% Normal Goat Serum (m/v) (S26-LITER, Merck Life Science UK LTD) (PBST.NGS) for 3 × 2 hours on an orbital shaker (160 rpm) at room temperature. Ears were then gently shaken (120 rpm) at 4°C overnight in 20 μg/ml Dylight 488 strepavidin (SA-5488, Vector Laboratories) and 0.05 mg/ml DAPI (D9542, Sigma Aldrich), diluted in PBST.NGS. During this time the fluorescent strepavidin binds very tightly to the fixed neurobiotin to specifically stain the recorded neurons and DAPI specifically binds to cell nuclei. After this overnight incubation, Müller’s organs were washed three times in PBS (3×10 min), dehydrated in an ethanol series and cleared in Methyl salicylate (M6752, Sigma Aldrich).

Fluorescence images (pixel size 0.31 μm^2^, with a total of 65 z stacks) were captured with a confocal microscope (FV1000 CLSM, Olympus) equipped with Plan-UPlanSApo 20x (0.75 numerical aperture) lens. Fluorescence emission of Dylight 488 was collected through a 505-530 nm bandpass filter and fluorescence emission of DAPI was collected through a 485 nm low pass filter. Confocal images were adjusted for contrast and brightness, overlaid and stacked in ImageJ (version 1.51, National Institutes of Health).

### Morphological analysis and cell counting of Müller’s organ

To count the auditory neurons, a maximum intensity z-projection was produced for each z-stack image in ImageJ, displaying the auditory neurons as a 2D image. Neurons were manually counted using the Cell Counter plugin (Kurt de Vos, University of Sheffield, https://imagej.nih.gov/ij/plugins/cell-counter.html). Contrast/ brightness parameters were adjusted to ensure all neurons were counted.

Imaris software was used for automated cell counts of the DAPI stained nuclei present in Müller’s organ and nerve. Automated cell counting was performed on 3D images using the “Spots” tool, with consistent parameters (15 μm spot size) used across all Müller’s organs. The “Surfaces” tool was used to ensure only cells within the Müller’s organ were included in the automated counting. The width of the auditory nerve was also measured in Imaris using the MeasurementPro tool.

### Experimental design and statistical analysis

We designed all experiments to have a power above 95%, which gives a false positive rate (probability of finding an effect that is not there) of 5%. In order to calculate the power we used the raw data, and the effect size, reported in Warren et al. (2020) for: Tympanal displacements measured with Doppler laser vibrometry (Figure 1Ai), hook electrode recordings of tone-evoked potentials from the auditory nerve (Figure 2) and whole-cell patch-clamp recordings from individual auditory neurons of Müller’s organ (Figure 3I). There exists no analytical methodology for conducting power calculations on Linear Mixed Effect Models (LMEM). Therefore we generated a dataset simulated from the raw data of Warren et al. (2020), fitted a Linear Mixed Effect Model, then ran repeated simulations of the LMEM 1000 times. We used the proportion of time that the LMEM reported a difference to calculate the power. For this paper the experiments that measured differences in tympanal displacements, hook electrode responses and the transduction current all had at least 95% power when using n numbers of 16, 9 and 12 respectively. This simulated power analysis also, only applies for the effect size reported in Warren et al (2020) and may be lower if the actual effect size, in this paper, is reduced. This is especially true for tympanal displacement measurements which were only ∼twice as different in these recordings as opposed to ∼10 times different in Warren et al. (2020). Models were fitted in R (Version 2.4.3), on a Windows PC running Windows 10 using the package *LME4* (Bates et al., 2015) and simulations were run with the package *simr* (Green and Macleod, 2016).

**Figure 2.**
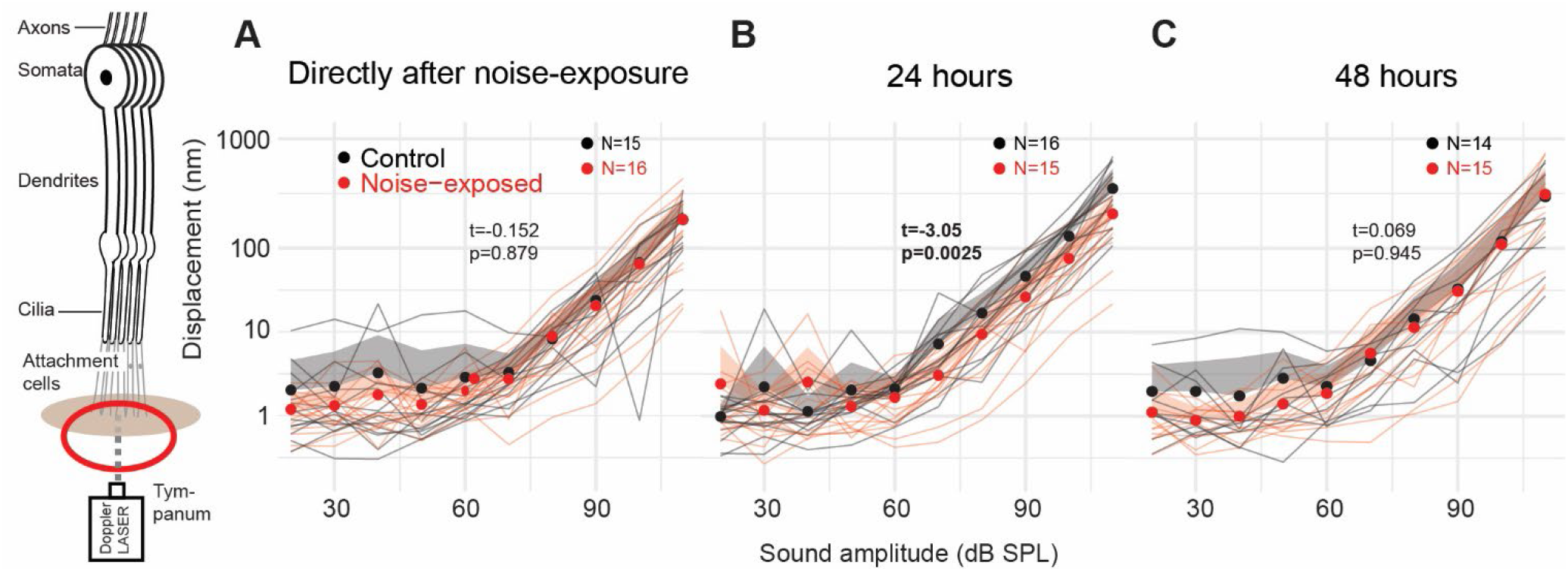
Doppler laser measurements of tone-evoked tympanal displacements. **A** Displacements of the tympanum (measured at the foot of the styliform body) in response to 3 kHz pure tones for control locusts and for locusts directly after noise exposure. Individual locusts are plotted as thin grey or thin red lines for control and noise-exposed respectively. Dots are the average for each treatment at each SPL. The positive standard deviation (negative not shown) is displayed in grey and red shaded areas. The extent of the difference between each group and a statistical test of a difference are displayed as t and p values.

**Figure 3.**
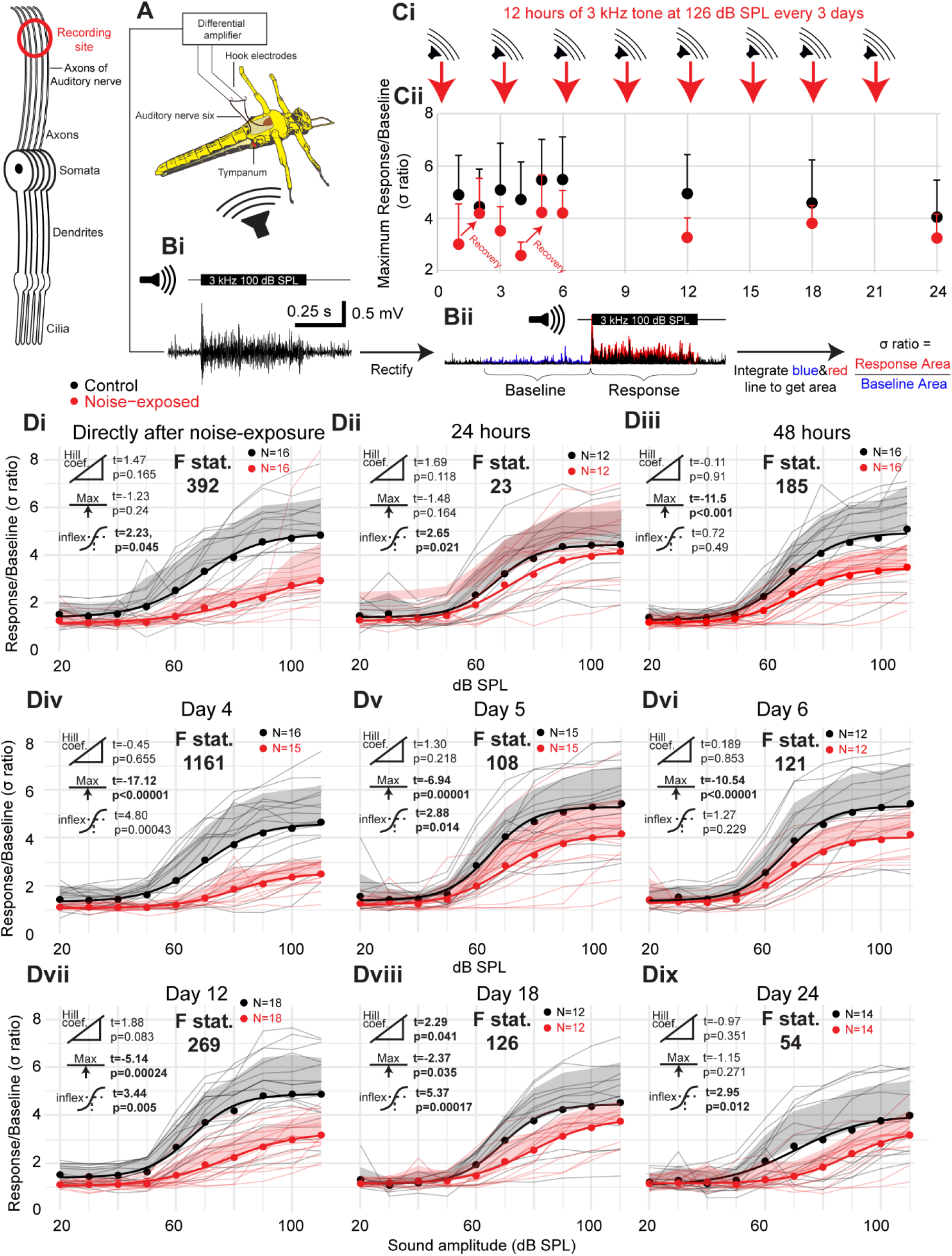
Hook electrode recordings from the auditory nerve quantify noise-induced and age-related hearing loss. **A** Schematic of dissection and recording setup. **Bi** Stimulation of 3 kHz pure tone elicits tone-evoked biphasic compound spikes. **Bii** Analytical workflow of tone-evoked responses. Auditory nerve signals were rectified and then the area underneath the tone-evoked signal was divided by the equivalent time and area with no auditory stimulation. **C** Experimental workflow showing the time of auditory over-exposures and recordings (N.B. days 1 and 4 where directly after noise-exposure and days 2 and 5 24hrs after noise exposure. Recordings on days 3, 6, 12, 18, 24 where all 48 hrs after noise-exposure and directly before the next noise exposure). **Cii** Plot of the maximal s ratio for control and noise exposed locusts at each recording day with positive standard deviation plotted. **Di** Plot of the s ratio (nerve response) as a function of SPL of a kHz tone for control and noise exposed locusts within 12 hours after cessation of noise exposure. Responses for individual locusts are plotted as thin grey or thin red lines for control and noise-exposed respectively. Dots are the average for each treatment at each SPL, shaded area is the positive standard deviation at each SPL and thick lines are four-parameter Log-Linear fits for each treatment. The extent of the difference between each group and a statistical test of a difference are displayed as t and p values next to three icons donating (from top to bottom) Hill coefficient, maximum asymptote and the inflexion point (p values below 0.05 are in bold). The F statistic compares the fit of the model when the treatment of the group (noise-exposed or control) was omitted. Higher F statistics donate a larger effect of treatment. **Dii** and **Diii** plot s ratio against SPL for noise-exposed and control locusts 24 hrs and 48 hrs after cessation of noise exposure (or silent speaker for control). **Div** is directly after a second noise exposure and **Dv** and **Dvi** 24hrs and 48hrs after the second noise exposure. **Dvii, Dviii**, and **Dix** are 48 hours after the 4^th^, 6^th^ and 8^th^ noise exposure respectively.

To calculate the number of locusts required to maintain a power of 95% for Müller’s organ morphological analysis we used preliminary data collected by Georgina Fenton. Georgina found an effect size *d* of 1.45 between the number of cells in Müller’s organ at 10 days old and 38 days old (two independent groups). We used G*Power t-test A Priori power analysis to calculate that a total sample size of 24. We settled on a sample size of 20, which still achieves a power of at least 95% when regression analysis across all five time points are taken into account (e.g. at days: 0, 6, 12, 18, 24 Fig. 1B).

Throughout the manuscript n refers to the number of recorded neurons and N refers to the number of Müller’s Organ preparations used to achieve these recordings (i.e. n=10, N=6 means that 10 neurons were recorded from 6 Müller’s Organs). All n numbers are displayed on the figures for clarity. The Spread of the data is indicated by 1 standard deviation as the standard deviation indicates the spread of the data, unlike standard error. Median and Q1 and Q3 are displayed by bars when individual measurements are plotted. For all hook electrode recordings, 80% of patch-clamp recordings and 50% of neurobiotin backfills the treatment of the locust (noise-exposed or control) was blinded to the experimenter; lone working conditions, due to Covid restrictions, made complete blinding impossible. All data either remained blinded or was recoded to be completely blind when analysing the data to avoid unconscious bias.

To test for differences and interactions between control, noise-exposed and aged locusts we used either a linear model (LM) or Linear Mixed Effects Model (LMEM), with treatment and age as fixed effects, and Locust ID and SPL as a random intercept, when repeated measurements are reported. Models were fitted in R (Version 3.4.3) with the package *LME4* (Bates et al., 2015). The test statistic for these analyses (t) are reported with the degrees of freedom (in subscript) and p value, which are approximated using Satterthwaite equation (lmerTest package) (Kuznetsova et al., 2017). We report Cohen’s d effect size for significant differences. Curves where fitted to the data using Matlab (version R2018a) for hook electrode recordings or the *drm* package in R for patch-clamp recordings (Ritz, 2016). The *drm* package was also used to compute t and p values when comparing control and noise-exposed four part Log-Linear models. F statistics of the Log-Linear model fits were computed by excluding treatment (noise-exposed or control) as a factor. Higher F statistics donate a stronger effect of treatment.

In order to compare responses between noise-exposed and control locusts across SPLs we adopted an approach first implemented in pharmacology research. In our work the “dose-response curves” are equivalent to SPL-auditory response curves. This allowed us to maximise the information contained in each dataset and to quantitatively compare model parameters such as: Hill coefficient (steepness of slope), maximal asymptote (maximum σ ratio), and inflexion point (σ ratio at the steepest part of the slope). We did this using the *drm* function of *drc* package (Version 3.1-1, Ritz and Strebig, 2015). We fitted four-part Log-Linear models with auditory nerve responses (σ ratio) as the dependent variable with treatment (control or noise-exposed) and SPL as the independent variables. This analysis was done for each day and the t and p value reported for each model parameter: Hill coefficient, maximal asymptote and inflexion point on each graph in Figures 2 and 3.

To test whether the factor of treatment (noise-exposed or control) significantly affected auditory nerve response we compared the above model to a model in which treatment was omitted as an independent variable, using the *anova* function (Ritz et al., 2015). This gave an F statistic labelled on each graph in Figures 2 and 3. The p value for this analysis remained below 0.001 for all graphs. We used identical analysis for whole-cell patch-clamp data where transduction current was the dependant variable.

Analysed datasets are freely available to download: https://www2.le.ac.uk/departments/npb/people/bw120. Please let me know if you require the raw data and especially if you wish to do a quantitative analysis of sex differences in hearing loss.

## Results

### Doppler Laser Vibrometry

Charlie Woodrow and Alix Blockley measured *in vivo* tone-evoked displacements from the external side of the tympanum at the foot of the styliform body, where the majority of group-III auditory neurons attach, using contactless laser Doppler vibrometry. Tone-evoked displacements rose above the noise floor above sound amplitudes of 60 dB SPL (Fig. 2A-C) and higher SPLs remained linear on a log-log plot. Thus, tone-evoked tympanal displacements can be explained by a simple power law. Directly after, and at 48 hours after noise exposure, there was no difference in tone-evoked tympanal displacements for control and noise exposed locusts (LMEM with SPL as a random effect; t_(297)_=-0.152, p=0.879; t_(209)_=0.069, p=0.945). The tympanal displacements of the noise-exposed locusts 24 hours after noise exposure were higher (t_(298)_=-3.05, p=0.0025).

### In vivo electrophysiology

Alix Blockley quantified the *in vivo* performance of the auditory neurons to produce sound-evoked spikes in response to repeated noise exposure over an extended 24 day period. She recorded spiking activity directly from the auditory nerve (nerve 6) using hook electrodes in response to 3 kHz tones produced by a speaker (Fig. 3A, B). A 3 kHz tone matched the frequency of our noise exposure and the frequency to which the majority of auditory neurons (Group III neurons) in Müller’s organ are broadly tuned. These recordings left the bilateral ears, in the first abdominal segment, intact (Fig. 3A). She exposed locusts to a 126 dB SPL 3 kHz tone overnight for 12 hours every three days and recorded nerve activity at days: 1,2,3,4,5,6,12,18,24 (Fig. 3 C).

The tone-evoked activity of the auditory nerve were composed of biphasic spikes that summated together at tone onset and for loud SPLs. Therefore, to quantify the tone-evoked activity of the auditory nerve she rectified the tone-evoked signals, integrated the area underneath and averaged the three tone presentations at each SPL. She then computed the σ ratio by dividing the average area during the 0.5 s tone by the average area over 0.5s during 60 seconds of background activity before tone presentations began (Fig. 3Bii). This normalises for differences in background spiking activity and noise of the recording. To maximise the information from the recordings we fitted four part Log-Linear equations to model the relationship between transduction current and SPL (Fig. 3Di-Dix). We compared three specific parameters of four part Log-Linear equations between the nerve response of control and noise exposed locusts: Hill coefficient, maximum asymptote, sound pressure level at inflexion point; and one general fit, the F statistic. Directly after the first noise exposure the spiking activity of the auditory nerve is decreased, compared to controls when taking into account the dependence of sound pressure level on the transduction current (Locust condition * SPL interaction) (Fig. 3Di, LMEM t_(315)_= -6.60 p<0.000001, Cohen’s effect size d (at 90 dB SPL)= -2.04, **F statistic = 392**), but recovers within 24 hours (Fig. 3Dii, LMEM t_(235)_= -0.910 p<0.364, **F statistic = 23**) before once again becoming more distinct 48 hours afterwards (Fig. 3Diii, LMEM t_(316)_=5.2, p<0.00000035, **F statistic = 185**). Directly after the second noise exposure (Day4) the spiking activity of the auditory nerve is decreased (Fig. 3Div, LMEM t_(305)_= -7.28 p<0.000001, Cohen’s effect size d (at 90 dB SPL)= -1.73. **F statistic = 1161**), and remains lower than controls on day 5 (Fig. 3Dv, LMEM t_(295)_= -3.56 p=0.000431, **F statistic = 108**) and day 6 (Fig. 3Dvi, LMEM t_(235)_=-3.72 p=0.000253, Cohen’s effect size d (at 90 dB SPL)= -0.99, **F statistic = 121**). On day 12 after four 12 hr noise-exposures the decrease in the sound-evoked nerve response is most severe (Fig. 3Dvii, LMEM t_(355)_= -6.49 p<0.000001, Cohen’s effect size d (at 90 dB SPL)= -1.73, **F statistic = 269**) before age-related hearing loss starts to narrow the difference between control-aged locusts and noise-exposed-aged locusts (Fig. 3viii, ix: Day 18 LMEM t_(235)_= -3.36 p<0.0009, Cohen’s effect size d (at 90 dB SPL)= -0.90 **F statistic = 126**; Day 24 LMEM t=_(275)_-3.33 p<0.00097, Cohen’s effect size d (at 90 dB SPL)= -0.750, **F statistic = 66**)).

### Morphology and number of auditory neurons and supporting cells in Müllers organ

Daisy Ogle backfilled the auditory nerves of control locusts and noise-exposed locusts *in vivo* with neurobiotin. She then stained auditory nerves with strepavidin florescent dye and the nuclei of all cells with DAPI. She counted the number of auditory neurons and number of cells in Müller’s organ, the density of cells in the auditory nerve and auditory nerve width. The auditory nerve width was reduced after noise exposure (Fig. 4A, B) (Linear Model: t_(61)_= -2.38, p= 0.0204) but did not decrease as a function of age (Linear model: t_(61)_= -0.675, p=0.502). There was no profound decrease in the number of cells in the auditory nerve as a function of age (LM: t_(61)_= -1.19, p=0.239) and noise exposure (LM: t_(61)_= -1.305, p= 0.198). Similarly, the number of cells in Müllers organ did not change as a function of age (LM: t_(77)_= -0.313, p= 0.755) or noise exposure (LM: t_(77)_= 0.419, p= 0.676). The number of auditory neurons strongly decreased as a function of age (LM: t_(77)_=-8.10 p<0.00001) and a small compounding effect of noise exposure of the number of auditory neurons increased as a function of age (day6, t_(10)_= -0.49, day12, t_(17)_= -0.91, day18, t_(16)_= -1.10, day24, t_(14)_= -1.89).

**Figure 4.**
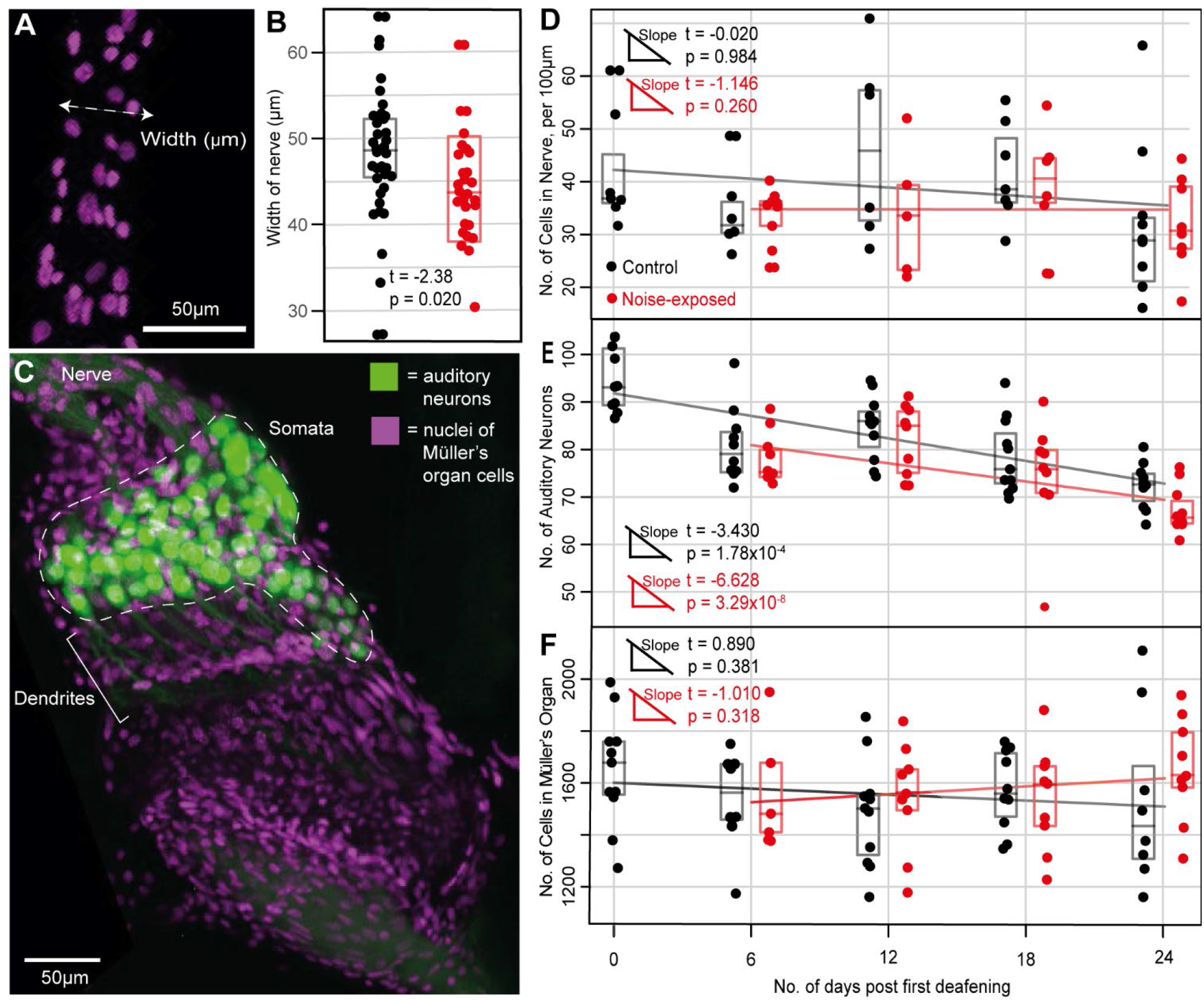
Anatomical analysis of Müller’s organ for noise-exposed and control locusts. **A** DAPI (nuceli) staining of cells in auditory nerve showing measurement of nerve width. **B** Comparison of auditory nerve width for noise-exposed and control locusts. **C** Dual staining of auditory neurons and DAPI staining of the nuclei of all cells in Müller’s organ. **D** Number of cells per 100 μm in the auditory nerve in noise-exposed and control locusts over 24 days. Linear regression analysis output is plotted for each treatment group. **E** Number of auditory neurons in noise-exposed and control locusts’ ears over 24 days. **F** Number of cells in Müller’s organ in noise-exposed and control locusts over 24 days.

### Electrical properties of auditory neurons and transduction current dependence on sound pressure level

We measured the electrical properties and tone-elicited currents of Group III neurons in Müller’s organ through whole-cell patch-clamp recordings from individual neurons from excised ears (Fig. 5A). We isolated and optimised the transduction current at the distal ciliated end of the auditory neuron using pharmacology, voltage protocols and the optimal sound stimulus (Fig. 5B) (detailed in methods). We recorded directly after the first 12 hour noise exposure at day 1 then 24 hours and 48 hours afterwards on day 2 and 3. Recordings at day 6, 12, 18, 24 were all made 48 hours after the last 12 hour noise exposure (Fig. 5C). The transduction current increased with an increased sound amplitude and followed a log-linear model (Fig. 5D). Directly after noise-exposure the maximum transduction current is reduced (Fig. 4Di, t=_(17)_-3.64, p=0.0018, Cohen’s effect size d (at 90 dB SPL)= -1.03) and reduces further 24 hours after noise exposure (Fig. 5Dii, t=_(17)_-8.08, p<0.00001 Cohen’s effect size d (at 100 dB SPL)= -1.28). Only 48 hours after the first noise exposure is the maximum transduction current comparable with non-noise-exposed controls. Interestingly the Hill coefficient, which mathematically derives the interaction between sound pressure level and transduction current, does not change directly after noise-exposure (Fig. 5Di t=_(17)_1.66, p=0.113), but changed after 24 hours (Fig.5Dii t=3.76, p=0.0014 and remains different 48 hours later (Fig. 5Diii, t=_(17)_3.87, p=0.0011). After two noise exposures and a 48 hour recovery (Day 6) both groups are still fairly similar (Fig. 5Div). Upon further noise-exposure there is further decline in the maximum transduction current of the noise-exposed group (Fig. 5C, Dv-Dvii Cohen’s effect size d (at 110 dB SPL)= -1.24) with the F statistic at its highest (492) when comparing control and noise-exposed locusts at 24 days whereas the control locusts show no age-related decline (Fig. 5C, Dv-Dvii).

**Figure 5.**
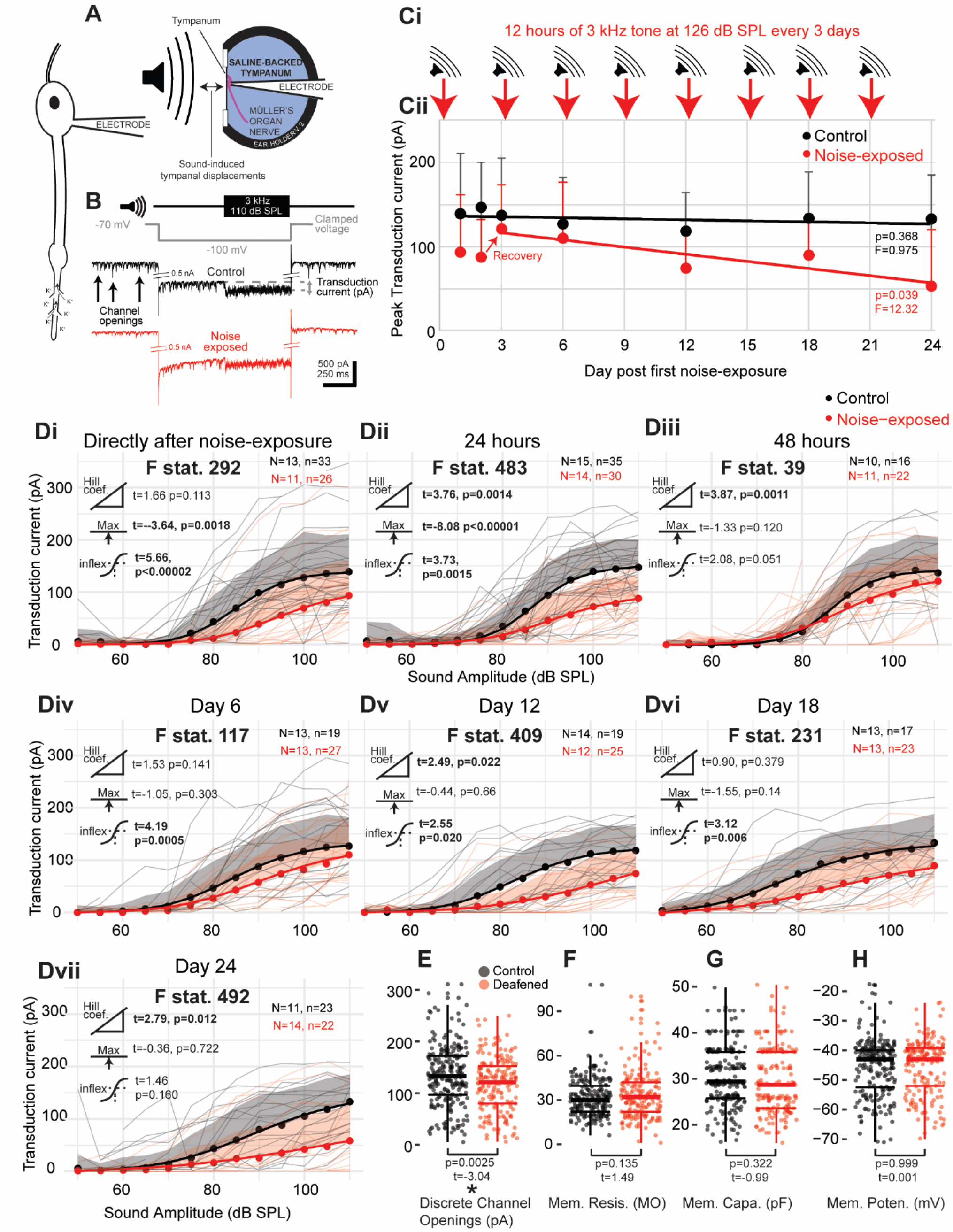
Analysis of tone-evoked transduction currents and electrophysiological properties for individual auditory neurons from noise-exposed and control locust ears. **A** Schematic depicting experimental setup. **B** Example of recording of the transduction current, including tone and voltage clamping protocols. **Ci** Experimental workflow of noise-exposure and mock noise exposure (for control locusts). **Cii** Plot of average maximum transduction current for noise-exposed and control locusts for each day. **Di** Plot of the transduction current as a function of SPL of a 3 kHz tone for control and noise exposed locusts within 12 hours after cessation of noise exposure. Transduction currents for individual locusts are plotted as thin grey or thin red lines for control and noise-exposed respectively. Dots are the average transduction current for each treatment at each SPL and shaded region is positive standard deviation for each treatment and SPL. Thick lines are four-parameter Log-Linear fits for each treatment. The extent of the difference between each group and a statistical test of a difference are displayed as t and p values next to three icons donating (from top to bottom) Hill coefficient, maximum asymptote and the inflexion point. Values are in bold when p<0.05. The F statistic compares the fit of the model when the treatment of the group (noise-exposed or control) was omitted. Higher F statistics donate a larger effect of treatment. **Dii** and **Diii** plot transduction current against SPL for noise-exposed and control locusts 24 hrs and 48 hrs cessation of noise exposure (or silent speaker for control). **Div, Dv, Dvi** and **Dvii** are 48 hours after the 2^nd^, 4^th^, 6^th^ and 8^th^ noise exposure respectively. **E** Plots the average discrete depolarisation amplitude for each auditory neuron for auditory neurons from noise-exposed and control locusts. The extent of the difference and the statistical test for any difference are plotted as t and p values respectively. **F, G, H** Plots the membrane resistance, membrane capacitance and membrane potential for each auditory neuron from noise-exposed and control locusts. Linear models were used to test for differences for electrophysiological properties of auditory neurons from control and noise-exposed locusts, with their t and p values displayed on the figures.

We recorded smaller discrete depolarisations in the auditory neurons of noise-exposed locust auditory neurons compared to control (Fig. 5E, 3E; LM t_(413)_=-3.104, p<0.002, Cohen’s effect size d=-0.306). Discrete depolarisations are assumed to be transient stochastic opening of mechanotransduction ion channels shown both in insects (Hill, 1983b; Warren & Matheson, 2018) and vertebrate auditory receptors (Beurg et al., 2015; Pan et al., 2013). There were no differences in the membrane resistance, membrane capacitance and membrane potential (at zero holding current) for the auditory neurons of control and noise-exposed locusts (Fig. 5F, G, H). However, there was a trend for the resting potential of auditory neurons of aged locusts (regardless of noise-exposure) to be ∼3 mV depolarised over 24 days of aging (LM; t=2.382, p=0.0177).

## Discussion

Repeated or prolonged noise exposure causes hearing loss which, for humans, is most different from non-noise-exposed individuals during the middle of their life (Passchier-Vermeer, 1974). As humans age, age-related hearing loss dominates and noise-exposure plays less of a role in determining hearing loss (Corso, 1980; Echt et al., 1998). The interaction of age and noise on hearing loss is only known for the human auditory system thanks to extensive longitudinal audiometric measurements including from industrial workers before ear protection (Passchier-vermeer, 1968; Corso, 1963). Here, we repeatedly exposed large numbers of locusts to noise and quantified their ability to hear over their life by measuring the response of their auditory nerve to sound. We also measured components of the auditory system known to correlate with hearing loss: decrease in auditory receptors, auditory nerve thinning, decrease of electrochemical gradients and other components crucial to the function of the auditory system: supporting cells, auditory receptor electrophysiological properties and transduction. We found striking parallels with the pattern of human hearing loss and rigorously quantify – for the first time in any auditory system – how noise and age affect specific components in the ear.

Before sound can impinge on the delicate inner ear sensory structures it must be captured mechanically through a tympanum (for sound pressure changes) or antennae (for particle velocity changes). Although there are clear age-related anatomical changes in middle ear bones and the tympanum in mammals this leads to no consistent changes in middle ear impedance, even in studies with different sexes and races (Thompson et al., 1979; Wada and Koike, 1994; Holte, 1996). There is a lack of studies on the effect of noise exposure (outside of blast type injuries (Gan et al., 2016)) on middle ear mechanics but, in flies, noise exposure increases the sharpness of tuning of its antennal sound receiver due to increased metabolic activity, and motile output, of the sensory auditory neurons (Boyd-Gibbins et al., 2021). Noise-associated changes in the locust ear tympanum are severe by comparison with any other auditory system with an increase of ∼10 times in its displacement after a 24 hour noise exposure (Warren et al., 2020). Here, with only a 12 hour noise exposure, there was only mild and transient decrease in tympanal displacements 24 hours after noise exposure. Thus, noise < 12 hours (at 126 dB SPL) appears to have a small, delayed and transient effect on the biomechanics of the locust’s tympanum.

We measured compound spiking potentials from the auditory nerve in response to a 3 kHz pure tone at SPLs from 20-110 dB SPL. We found a decrease in maximal nerve response directly after noise exposure (Fig. 3Di, Div). This is similar to a decrease in the first wave of auditory brainstem responses (ABRs) and compound action potentials after noise exposure in mice (Kujawa and Liberman, 2009; Chuang et al., 2014) rats (Tagoe et al., 2014). Noise-exposed locusts recovered their maximal nerve response within 24 hours after first noise exposure but it then decreased again at 48 hours later (Fig. Di-Diii), a 48 hour pattern repeated after the second noise-exposure on day 4 (Div-Dvi). By comparison mice (with an accelerated hearing loss phenotype) recover quickest in the first three days before their recovery slows with no sign of a reversal apparent within the days measured (straight after, 1, 3, 7 and 14 days after noise exposure) (Chuang et a., 2014). And other studies that measured hearing loss after noise exposure have only ever found recovery (Li and Borg, 1993; Howgate and Plack, 2011; Chuang et al., 2014; Fernandez et al., 2015; Kujawa and Liberman, 2009). Although the reversal of recovery found for locusts appears paradoxical, no study has measured recovery twice within 48 hours to uncover short term changes. This pattern of recovery mirrors some traumatic brain injuries where initial recovery can be followed by worsening of symptoms. The physiological basis of such non-linear recovery is due to different aetiologies presenting over distinct time courses. In the case of TBI initial electrophysiological dysfunction is followed by structural damage then toxic-metabolic and vascular effects (Giza and Hovda, 2001). We suggest that noise exposure has a complex recovery process where an initial promising recovery is followed shortly after by deterioration and is the result of multiple processes, triggered by noise exposure, acting over distinct time courses. We do not yet know if this reversal of recovery after noise exposure is an exception compared to the ears of other animals exposed to noise.

The long-term effects of repeated noise exposure result in a decreased maximum nerve response and higher SPL at the inflexion point (indicative of increased hearing thresholds). But age-related hearing loss leads to narrowing of the differences between control and noise-exposed locusts at older ages. This follows human hearing loss – the only other organism with a high enough powered comparative longitudinal dataset (Corso, 1980) and suggests this pattern of degeneration is fundamental across diverse biological auditory systems.

Hair cell number decreases after loud or prolonged noise exposure (Dixon and Ward, 1971; Clark et al., 1974; Moody et al., 1976; Bohne and Harding, 2000; Chen and Fechter, 2003). The 126 dB SPL 3 kHz tone noise exposure, used here to deafen locusts, is at similar sound levels (to the above studies) but resulted in little effect on the loss of locust auditory receptors. Instead auditory receptor number decreased steadily due to age with a mild compounding effect of noise. Mild (100 dB SPL) noise exposure in mice appears to do little to compound a strongly age-related auditory receptor decline (Fernandez et al., 2015). If this is similar in humans then (we cautiously conclude) noise-induced hearing loss may have little to do with loss of hair cells – at least for physiologically realistic SPLs experienced by animals and most humans. The myelin sheath encapsulating auditory nerves of mammals is damaged after noise exposure (Pilati et al., 2012; Tagoe et al., 2014; Wan and Corfas, 2017) and auditory nerve thinning in locusts with noise suggests a similar consequence of noise exposure. The number of non-sensory cell types has not been counted in any auditory system and we found no decrease in non-neuronal cell types, either in the nerve or Müller’s organ, in the locusts either as a function of noise or age.

In individual auditory neurons of Müller’s organ, membrane resistance and cell capacitance were unaffected either by noise or age. There was a trend for older auditory neurons (irrespective of treatment) to be more depolarised ∼3 mV, although the magnitude of the change is small (t=-2.382, p=0.0177). Whilst there are no changes in the membrane resistance and membrane potential of hair cells of mice, their capacitance decreased as a function of age (Jeng et al., 2020). Properties of the transduction current in mice were unaffected by age which matches that found here for auditory neurons of locusts. This reveals remarkable homeostatic mechanisms at play to maintain auditory receptor function. Repeated exposure to noise, however, results in a steady decrease in the electrochemical driving force for the transduction current. In mammals the electrochemical driving force for the scala media is established by specialised ion-pumping cells in the stria vascularis. Dysfunction of the stria vascularis has been recently popularised as responsible for the majority of age-related hearing loss (Ramadan and Schuknecht, 1989; Vaden et al., 2017). More recent interpretations (Wu et al., 2020) suggest that although the correlation of stria vascularis atrophy in patients with hearing loss is high (Schuknecht, 1974; Schuknecht, 1993) it does little to predict the extent of hearing loss.

Despite the clear anatomical differences in auditory systems of insects and vertebrates they share equivalent components: transduction on ciliated endings, electrochemical gradient across the auditory receptors and spiking axons that carry auditory information to the central nervous system. In addition, they share developmentally interchangeable transcription factors for ear development (Wang et al., 2002) and evolved from the same ancestral ciliated sensory appendage (Nowotny and Warren, 2020). As such, insects are a valuable resource to derive general principles of how auditory systems are affected by age and noise. This is keenly evidenced by the similarities in the pattern, causes and consequences of hearing loss which stand in stark contrast to what can be “misleading” anatomical differences.

## Acknowledgements

BW was funded by the Royal Society University Research Fellowship. AB and DO were funded by a Royal Society Enhancement Award. Work carried out by CW and FM-Z was funded by the European Research Council, Grant ERCCoG-2017-773067 to FM-Z. We acknowledge the help of Georgina Fenton and Megan Barnes for collecting preliminary data necessary to determine the length and sound amplitude of the deafening tone that produced a temporary hearing loss. We thank Tom Austin for help with hook electrode recordings. We acknowledge the help of James Blockley for help with analysis of hook electrode and laser data. We acknowledge the help of the Kees Straatman and the Advanced Imaging Facility at the University of Leicester. We would also like to thank Beth Fraser and Neil Rimmer for locust husbandry and Ben Cooper and Brendan O’Connor for help with statistical analysis. We would like to thank the following for help with blinding the treatment of the locust to the experimenter: Ben Cooper, Brendan O’Connor, Will Norton, Tom Matheson, Swidbert Ott and Zoe Bailey.

## Author contributions

Charlie Woodrow and Alix Blockley collected and analysed the Doppler laser data (Fig. 2). Alix Blockley and analysed the hook electrode data (Fig 3.). Daisy Ogle collected and analysed all morphological data on Müller’s organs and composed Figure 4. Alix together with Ben Warren conceived the experimental workflow of the paper. BW conceived the idea for the paper, collected and analysed the patch-clamp electrophysiological data and composed Figure 5, wrote the paper and performed all statistical analyses. All authors helped refine figures, write the paper and interpret data.

